# Single-cell analysis of the somatic mutational landscape in human chondrocytes during aging and in osteoarthritis

**DOI:** 10.1101/2024.08.24.609506

**Authors:** Peijun Ren, Chen Zheng, Yidan Pang, Shixiang Sun, Qiyang Wang, Wanxing Xu, Moonsook Lee, Yu Qiang, Zhenzhen Lu, Min Zhou, Jian He, Ningning Liu, Alexander Y. Maslov, Xiao Dong, Junjie Gao, Changqing Zhang, Jan Vijg

**Author notes:** These authors contributed equally to this work: Peijun Ren, Chen Zheng, Yidan Pang. Correspondence to: Prof. Jan Vijg, Center for Single-Cell Omics, School of Public Health, Shanghai Jiao Tong University School of Medicine, Shanghai 200025, China;. Department of Genetics, Albert Einstein College of Medicine, Bronx, NY 10461, USA; Prof. Changqing Zhang, Department of Orthopaedics, Shanghai Sixth People’s Hospital Affiliated to Shanghai Jiao Tong University School of Medicine, Shanghai, 200233, China; Dr. Junjie Gao, Department of Orthopaedics, Shanghai Sixth People’s Hospital Affiliated to Shanghai Jiao Tong University School of Medicine, Shanghai, 200233, China; Dr. Xiao Dong, Institute on the Biology of Aging and Metabolism, University of Minnesota, Minneapolis, MN 55455, USA;.

## Abstract

Somatic mutation is now recognized as a cause of multiple human diseases other than cancer. Osteoarthritis (OA), a highly prevalent age-related disease, has been associated with increased chromosomal abnormalities in articular cartilage of OA patients. Thus far no systematic attempt has been made to characterize the somatic mutational landscape of chondrocytes during normal aging and in affected cartilage of OA patients. Here we used single-cell whole genome sequencing to quantitatively analyze single-nucleotide variants (SNVs) and small insertions and deletions (InDels) in 100 single chondrocytes isolated from the cartilage of hip femoral heads of 17 subjects aged from 26 to 90 years, including 9 OA patients and 8 non-OA donors. Both SNVs and InDels were found to accumulate with age in chondrocytes with a clock-like mutational signature. Surprisingly, the age-related accumulation rate of these mutations was found to be lower in OA chondrocytes compared with chondrocytes from non-OA control subjects.

## Introduction

Somatic cells acquire and accumulate mutations throughout the life span due to errors in the repair and replication of DNA damage inflicted from a variety of sources, including hydrolysis and oxidation^1^. Somatic mutations cause cancer but have recently been shown to underlie a wide variety of human diseases other than cancer^2–6^. Somatic mutations are difficult to measure due to their random nature, which essentially constrains their detection through regular high-throughput sequencing, which does not allow distinguishing sequencing errors from true *de novo* mutations. However, recent innovations have led to new methods allowing accurate quantitative analysis of somatic mutations. One of these methods is single-cell whole genome sequencing after multiple displacement amplification, i.e., single-cell multiple displacement amplification (SCMDA)^7^.

Using SCMDA somatic mutations have been found to accumulate with age in human B cells, liver, and lung^8–10^. In lung, somatic mutation burden was found to be elevated in bronchial epithelial cells from smokers as compared to non-smokers. Using an alternative, accurate MDA procedure, i.e., linked read analysis, somatic mutations have also been shown to increase with age in human brain^4^ and heart^11^. In all these tissues single-nucleotide variants (SNVs) were found at numbers that varied from about 1,000 in cells from young subjects to several thousand per cell in people over 70.

Osteoarthritis (OA) is a major public health challenge world-wide. Globally, the age-standardized prevalence of OA shows an increase of 9.3% from 1990 to 2017^12^. The socio-economic burden of OA is increasing with population aging in most developed economies^13^. While advanced age is the primary risk factor for the disease^14^, the mechanisms underlying OA and how these may be linked to the aging process are currently unknown. Chondrocytes are the resident cells in the articular cartilage that secrete extracellular matrix and maintain the structure and balance of cartilage^15^. Hence, dysregulated chondrocyte physiology plays a key role in the initiation and progression of OA.

While OA has since long been associated with somatic mutations, mostly increased chromosomal abnormalities that could be readily detected^16,17^, thus far no attempt has been made using advanced single-cell sequencing methods to characterize the landscape of somatic mutations in chondrocytes during aging and OA. Here we fill this gap in our knowledge by applying single-cell whole-genome sequencing to individual chondrocytes from hip articular cartilage of donors with or without OA. We show that as in most other tissues that have been analyzed by advanced single cell sequencing methods, somatic mutations accumulate with age in human chondrocytes, revealing mutational signatures of normal aging. Surprisingly, somatic mutation burden in chondrocytes from OA patients were not higher than in the same cells from control subjects. Frequencies of both SNVs and InDels were significantly lower in OA chondrocytes than in non-OA control chondrocytes. In OA chondrocytes, elevated levels of apoptosis were observed, possibly due to the increased DNA damage which has been reported by others^18–20^. This may have led to the observed reduced somatic mutation accumulation by selectively eliminating damaged chondrocytes.

## Results

### Somatic mutation frequencies in chondrocytes increase with age

Somatic mutation burden was determined in articular chondrocytes isolated from cartilage in femoral heads (Fig. 1a; Methods). A total of 17 donors who received hip joint replacement surgery were recruited, aged between 26 and 90 years, including 9 hip osteoarthritis (OA) patients and 8 patients experiencing hip joint diseases other than OA (non-OA control) (Table 1). From 4 out of the 9 OA donors, we were able to collect cartilage samples from both a lesion site and a non-lesion site. From the remaining 5 OA donors the cartilage damage was severe, and no normal cartilage could be collected. The age range of the 8 non-OA donors (referred to as non-OA controls from now on) was 42 to 90 years. Individual cells were subjected to single-cell whole genome amplification and sequencing using SCMDA^7,21^. To subtract germline mutations, bulk genomic DNA for each subject was extracted from chondrocytes collected from the same cartilage samples and sequenced. In total, the whole genome of 100 single chondrocytes and 21 bulk DNAs was sequenced by paired-end sequencing using Illumina NovaSeq 6000 (Methods). According to cartilage damage status, the 100 chondrocytes were further divided into three groups: non-OA control (isolated from femoral head cartilages of non-OA donors), OA non-lesion (isolated from cartilages with normal physical appearance and morphology collected from femoral heads of OA donors) and OA lesions (isolated from damaged cartilage collected from femoral heads of OA donors).

**Figure 1.**
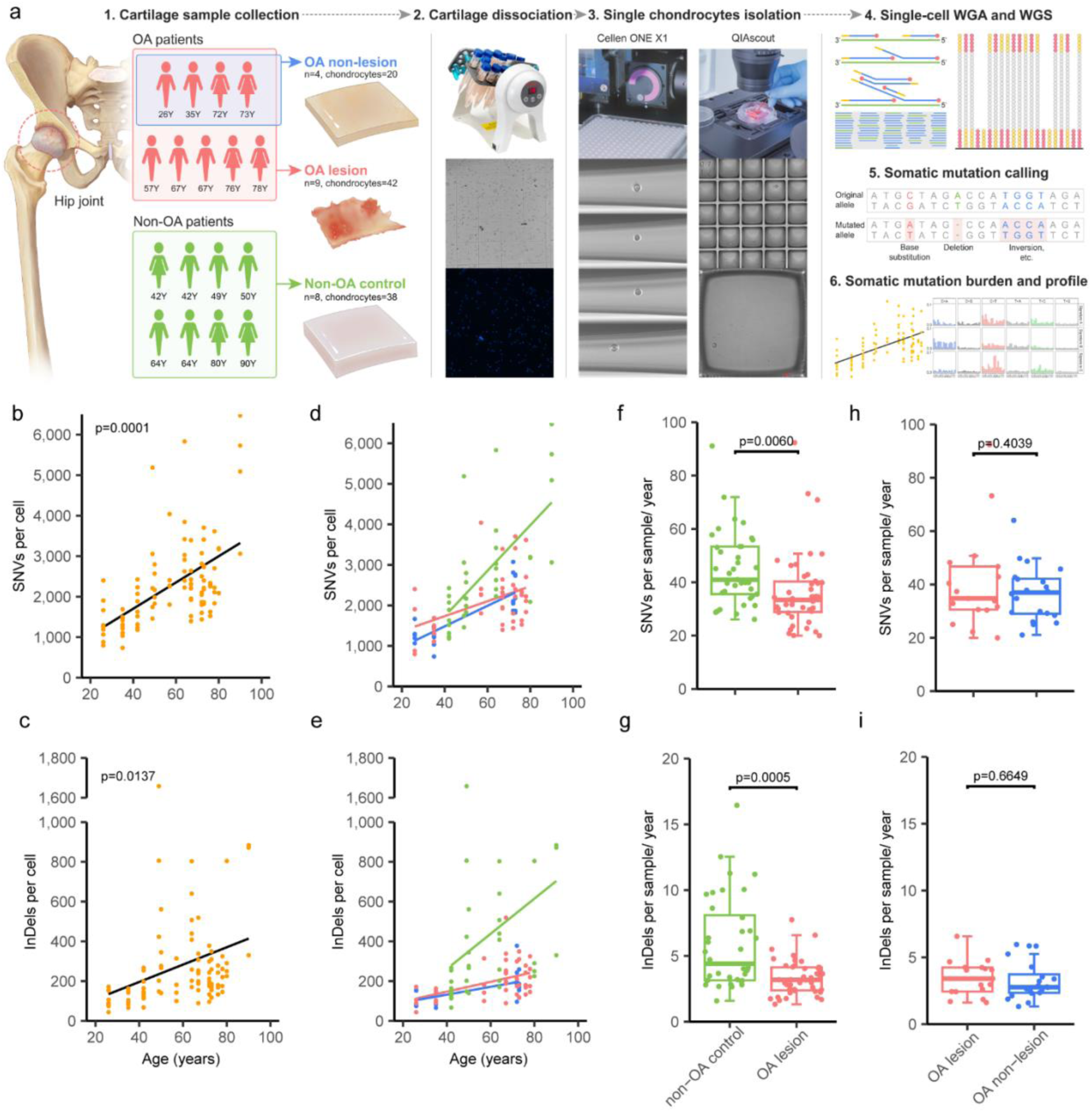
Somatic mutation accumulation in chondrocytes. **a,** Experimental workflow of single-cell WGS of chondrocytes from hip articular cartilage. Cartilage samples, collected from femoral head of donors who underwent arthroplasty, were classified into three groups: OA lesion (red), OA non-lesion (blue) and non-OA control (green). **b, c,** SNV and InDel frequency of all chondrocytes versus age, with linear mixed-effect regression lines. Each data point indicates the mutation frequency of individual chondrocytes (n=100). **d, e,** SNV and InDel frequency of chondrocytes versus age with linear mixed-effect regression lines in the non-OA control, OA lesion, and OA non-lesion groups. Each data point indicates the mutation frequency of individual chondrocytes. **f, g,** SNV and InDel accumulation rate of OA and control groups, with boxes indicating median values and interquartile range. Each data point indicates the per year mutation accumulation rate of each chondrocyte. **h, i,** SNV and InDel accumulation rate in chondrocytes of 4 paired cartilage samples, with boxes indicating median values and interquartile ranges, respectively. Each data point indicates the per year mutation accumulation rate of each chondrocyte. See also Supplementary Table 1.

**Table 1.** Case information and number of chondrocytes analyzed.

| Donor ID | Gender | Age | Diagnosis | Cartilage | Number of single cells |
| --- | --- | --- | --- | --- | --- |
| P1 | male | 64 | non-OA | control | 5 |
| P2 | male | 42 | non-OA | control | 5 |
| P3 | male | 64 | non-OA | control | 4 |
| P4 | male | 49 | non-OA | control | 5 |
| P6 | male | 50 | non-OA | control | 5 |
| P7 | male | 35 | OA | lesion | 5 |
|  |  |  |  | non-lesion | 5 |
| P8 | female | 42 | non-OA | control | 6 |
| P10 | female | 67 | OA | lesion | 5 |
| P11 | female | 67 | OA | lesion | 5 |
| P13 | male | 73 | OA | lesion | 4 |
|  |  |  |  | non-lesion | 5 |
| P15 | female | 72 | OA | lesion | 4 |
|  |  |  |  | non-lesion | 5 |
| P16 | male | 76 | OA | lesion | 5 |
| P19 | female | 78 | OA | lesion | 5 |
| P21 | male | 26 | OA | lesion | 4 |
|  |  |  |  | non-lesion | 5 |
| P22 | male | 57 | OA | lesion | 5 |
| P30 | female | 80 | non-OA | control | 4 |
| P32 | female | 90 | non-OA | control | 4 |

Among all sequenced single cells, the mean coverage was 35.20± 8.67% of the genome, which was surveyed at a depth of ≥20X. Single-nucleotide variants (SNVs) and small insertions and deletions (InDels) were detected using SCcaller, at a sensitivity for SNVs and InDels of 0.55 ± 0.10 and 0.29 ± 0.05, respectively, with germline variants filtered out (Methods and Supplementary Table 1). In all chondrocytes, the observed number of SNVs ranged from 135 to 835, with InDels varying between 3 and 83. After adjusting for genome coverage and sensitivity, the estimated number of SNVs per chondrocyte across all cells ranged from 737 to 5,830, and the number of InDels per chondrocyte was between 44 and 1,658 (Fig. 1b, c). As expected, the InDel frequency (mean: 283 per/cell) was significantly lower than the SNV frequency (mean: 2267 per/cell) (t-test, two-sided, p<0.0001). The linear mixed-effects model (LME) was used to analyze trends of SNVs and InDels with age. In chondrocytes tested, frequency of both SNVs and InDels increased linearly with age; SNVs accumulated at a rate of 32 per cell per year (LME, p=0.0001; Fig. 1b; Supplementary Table 1), while InDels accumulated at a rate of 4 per cell per year (LME, p=0.0137, Fig. 1c; Supplementary Table 1).

### Somatic mutation accumulation rate in OA chondrocytes is lower than that in non-OA subjects

To explore the pathogenesis of somatic mutations in OA, we compared age-related mutation accumulation in chondrocytes between OA subjects and non-OA controls (Fig. 1d, e). The mutation accumulation rate for each sample was then calculated by dividing mutation frequencies by age. The mean SNV accumulation rate in the non-OA controls was 47 SNVs/year, significantly higher than the 37 SNVs/year in the OA subjects (p=0.0060, t-test, two-sided, Fig. 1f). InDel accumulation rate in the non-OA control chondrocytes was 7 InDels/year, also significantly higher than in the OA chondrocytes which was 3 InDels/year (p=0.0005, t-test, two-sided, Fig. 1g).

Considering that for only 4 of the 9 OA donors matched lesion and non-lesion cartilage samples were available, data of these 4 paired samples were analyzed separately to validate whether cartilage damage affected somatic mutation profiles in chondrocytes. In the paired samples, SNV frequency showed no significant difference between the OA lesion group as compared to the OA non-lesion group (p=0.4039, two-sided, t-test, Fig. 1h); the same was true for InDel frequency (p=0.6649, two-sided, t-test, Fig. 1i).

### Mutational signatures distinguish OA from non-OA chondrocytes

We performed mutational signature analysis to identify specific mutational processes in chondrocytes during aging and OA. We extracted *de novo* mutational signatures from all SNVs and InDels detected in the 100 chondrocytes using non-negative matrix factorization (NMF) and observed 3 single base substitution signatures, termed signatures A, B and C (Fig. 2a, b, Extended data Fig. 1a), and 2 *de novo* InDel signatures, termed 1 and 2 (Fig. 3a, b, Extended data Fig. 1b)^22^.

**Figure 2.**
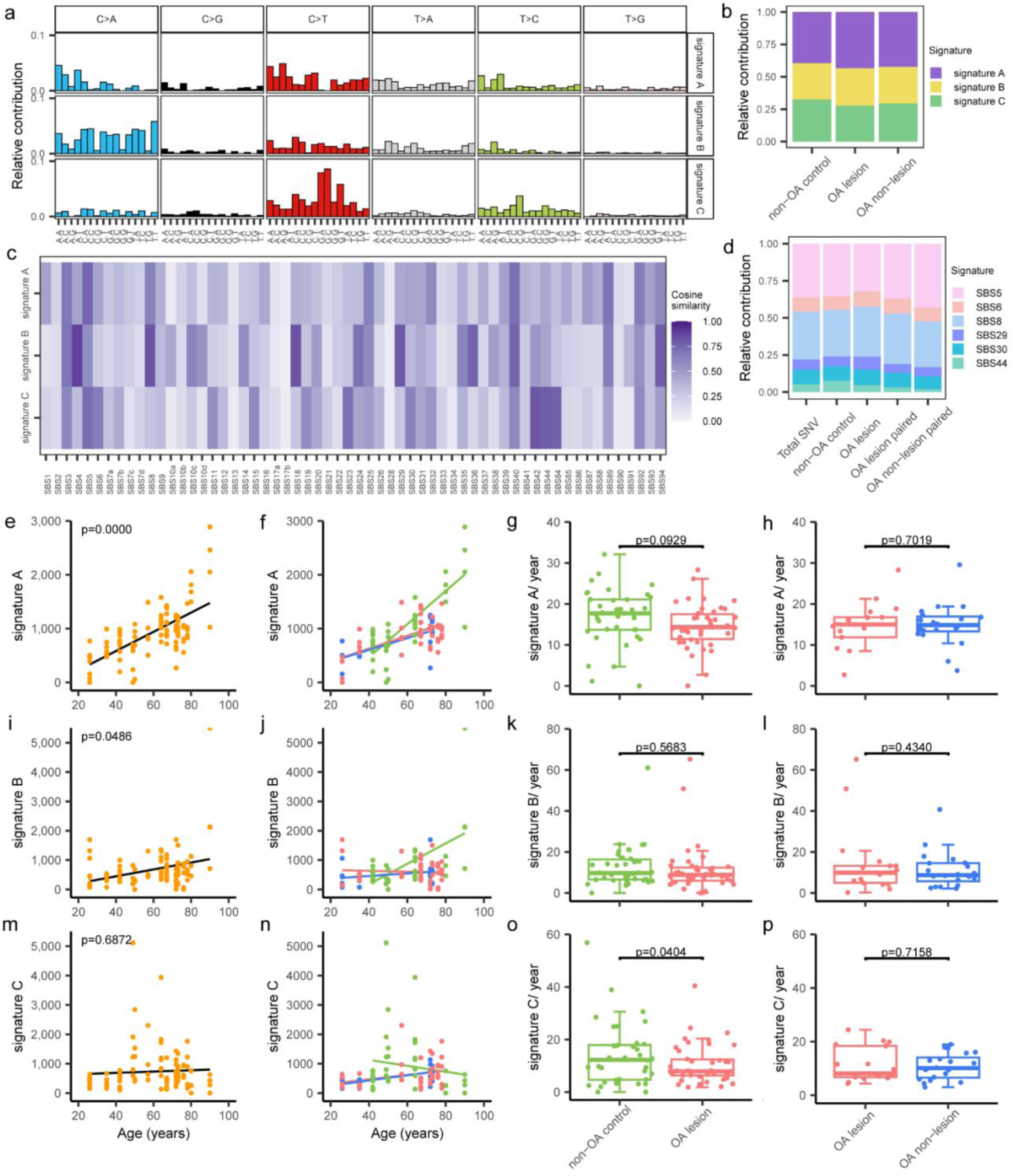
*De novo* signature analysis of single base substitutions in chondrocytes. **a,** Mutation spectra of three *de novo* single base substitution signatures extracted using NMF. **b,** The relative contribution of three signatures to all SNVs in three groups of cartilage damage status. **c,** Heat map showing the cosine similarities of signatures from COSMIC and extracted by NMF. Similarity coefficient is indicated by color scale at right. **d,** Signature refitting shows the relative contribution of COSMIC signatures in chondrocyte groups. **e, i, m,** Contribution of signatures to SNVs versus age in overall chondrocytes, with linear mixed-effect regression lines. Each data point indicates the mutation frequency per single chondrocyte. **f, j, n,** Contribution of signatures to SNVs versus age for each cartilage damage group, with linear mixed-effect regression lines. Each data point indicates the number of SNVs contributing to signature per single chondrocyte (n=100). **g, k, o,** Per year accumulation rates of signature contribution to SNVs for each single chondrocyte of different cartilage damage status groups, with boxes indicating median values and interquartile ranges of the non-OA control, OA lesion, respectively. Each data point indicates the per year mutation accumulation rate of each single chondrocyte contributing to the signature. **h, l, p,** Per year accumulation rates of signature contribution to SNVs for each single chondrocyte from paired cartilages, with boxes indicating median values and interquartile ranges. Each data point indicates the per year mutation accumulation rate of each single chondrocyte contributing to the signature. Group colors are as in Fig. 1.

**Figure 3.**
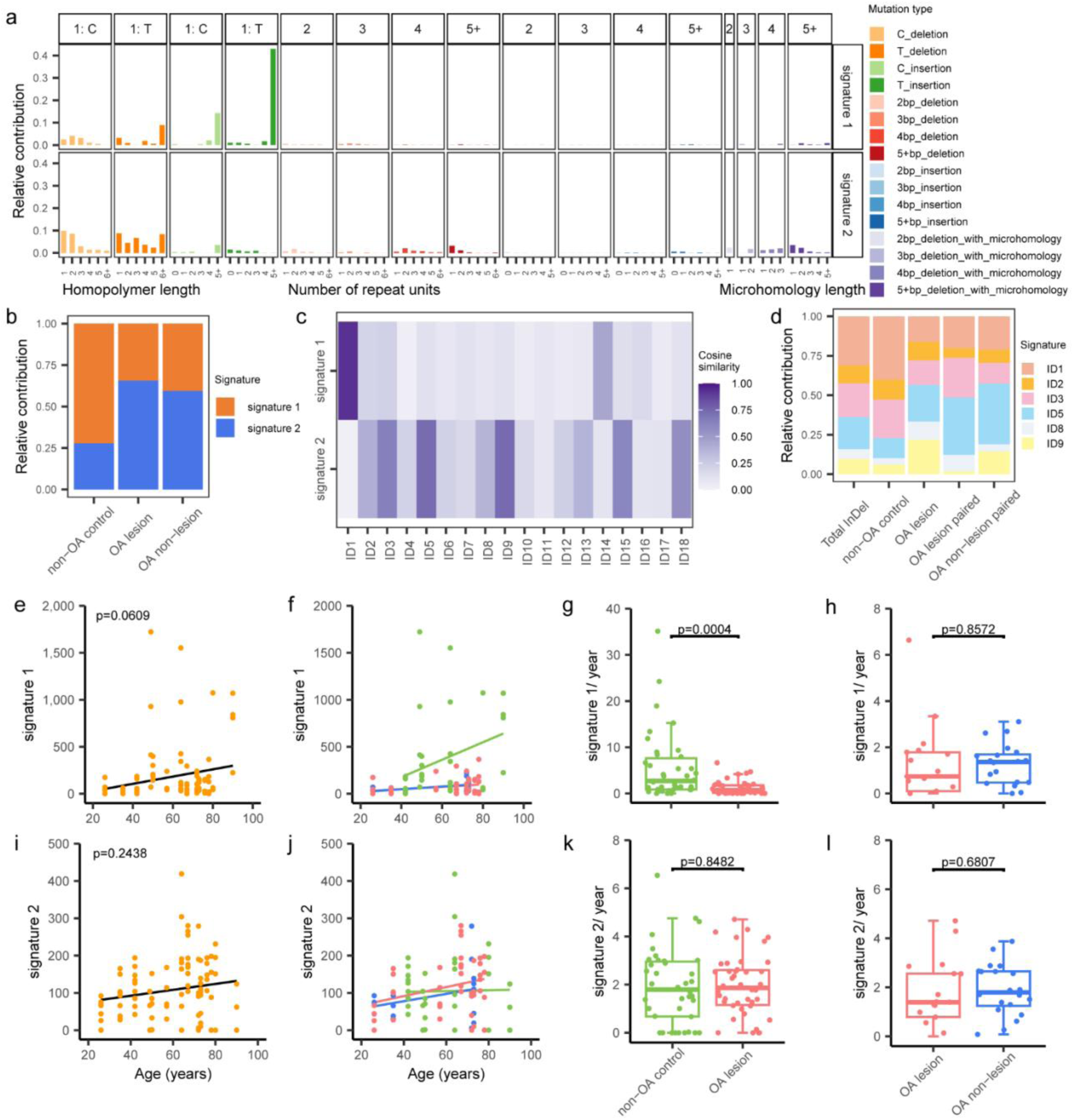
*De novo* signature analysis of InDels in chondrocytes. **a**, Mutation spectra of two *de novo* InDel signatures extracted using NMF. **b,** The relative contribution of two signatures to all InDels in three damage status groups. **c,** Heat map showing the cosine similarities of signatures from COSMIC and extracted by NMF. Similarity coefficient is indicated by color scale at right. **d,** Signature refitting shows the relative contribution of COSMIC signatures in chondrocyte groups. **e, i,** Contribution of signatures to InDels versus age in overall chondrocytes, with linear mixed-effect regression lines. Each data point indicates the mutation frequency per single chondrocyte. **f, j,** Contribution of signatures to InDels versus age for each cartilage damage group, with linear mixed-effect regression lines. Each data point indicates the number of InDels contribute to signature per single chondrocyte (n=100). **g, k,** Per year accumulation rates of signature contribution to InDels for each single chondrocyte of different cartilage damage status groups, with boxes indicating median values and interquartile ranges of the non-OA control and OA lesion, respectively. Each data point indicates the per year mutation accumulation rate of each single chondrocyte contributing to the signature. **h, l,** Per year accumulation rates of signature contribution to InDels for each single chondrocyte from paired cartilages, with boxes indicating median values and interquartile ranges. Each data point indicates the per year mutation accumulation rate of each single chondrocyte contributing to the signature. Group colors are as in Fig. 1.

Signature A was based mainly on C>T and T>C changes and had high similarity to SBS5 (cosine similarity of 0.76, Fig. 2c, Supplementary Table 2), a known age-related “clock-like” signature in the COSMIC mutational signature database (https://cancer.sanger.ac.uk/cosmic/signatures). SBS5 mutations linearly accumulate with age in all tissue types analyzed^23,24^. Fitting SNVs of signature A in linear mixed-effects models showed that its frequency increased significantly with age, at a rate of 18 SNVs/cell/year (p<0.0001, Fig. 2e, Supplementary Table 3). No significant difference of signature A accumulation rate was found between the OA and non-OA control groups (p=0.0929, Fig. 2f, g), nor among the paired samples (p=0.7019, Fig. 2h), indicating that the mutational processes forming signature A are independent from the disease-related cartilage damage.

Signature B was not found to correlate with age (Fig. 2i) and was also independent of cartilage damage status (Fig. 2j∼l). Signature B, which featured C>A substitutions with a smaller contribution of C>T and T>C mutations, was similar to SBS8 (cosine similarity: 0.82), SBS4 (cosine similarity: 0.88) and SBS18 (cosine similarity: 0.83, Fig. 2c). SBS8, which has been associated with deficiency in homologous recombination (HR) and nucleotide excision repair (NER)^24–26^, has been detected in neurons, cardiomyocytes and lymphocytes^4,11,27^. SBS4 has also been associated with NER, but more specifically with tobacco mutagens^28^. SBS18 has been associated with reactive oxygen species^23,29^.

The increase with age of signature C mutation frequency for OA and non-OA combined was not significant (p=0.6872, LME, Fig. 2m). When separating OA from non-OA, the accumulation rate in OA chondrocytes was slightly, but significantly lower compared with non-OA controls (p=0.0404, t-test, two-sided, Fig. 2n, o). There was no difference between paired lesion and non-lesion cartilages of OA donors (p=0.7158, t-test, two-sided, Fig. 2p). Signature C was similar to SBS42 (cosine similarity: 0.81) and SBS44 (cosine similarity: 0.79), recently been reported in cardiomyocytes. SBS44 has been associated with NER, but also with defective DNA mismatch repair (MMR)^11^.

For InDels, we detected two *de novo* signatures, termed signatures 1 and 2 (Fig. 3a, b, Supplementary Table 2 and 3). Neither signature 1 nor signature 2 was found correlated with age (Fig. 3e, i). The frequency and accumulation rate of signature 1 was higher in the non-OA control chondrocytes than in the OA chondrocytes (p=0.0004, t-test, two-sided, Fig. 3f, g), but showed no difference between paired lesion and non-lesion sites from OA cartilage (p=0.8572, t-test, two-sided, Fig. 3h). Signature 1 was close to ID1 (cosine similarity of 0.93, Fig. 3c) in COSMIC. ID1 is a signature common across tissues and has been associated with defective DNA mismatch repair (MMR)^24^. Signature 2 was independent of cartilage damage status (Fig. 2i∼l). Signature 2 was close to ID5 (cosine similarity of 0.74) and ID9 (cosine similarity of 0.73, Fig. 3c) for which the etiology is not known yet.

Next, we fitted known COSMIC signatures onto the 96 mutational spectra of all SNVs and 83 spectra for all InDels, which we detected in all 100 chondrocytes during aging and OA. In these aggregate mutation spectra, 6 single base substitution signatures contributed more than 4%: SBS5, 8, 30, 29, 44, and 6 (Fig. 2d, Supplementary Table 4). The clock-like signature SBS5 accumulated linearly with age in chondrocytes, with both frequency and accumulation rate independent from disease and cartilage damage status (Extended data Fig. 2a∼c, Supplementary Table 5), which is consistent with the *de novo* analysis above. For InDels, 6 signatures contributed more than 4%: ID1, 3, 5, 2, 9, and 8 (Fig. 3d, Supplementary Table 4). The proposed etiology of ID1 and 2 is slippage during DNA replication of the replicated DNA strand. ID3 has been associated with tobacco smoking.

### Genomic distribution of somatic mutations in chondrocytes

We annotated the distribution of all observed somatic mutations across the genome and observed significant mutation depletions in chondrocytes. Mutation depletion and enrichment analysis revealed significant SNV depletion from exon, promoter, and enhancer regions in chondrocytes from all three cartilage damage status groups (p<0.0001, Fig. 4a), confirming the well-documented effect of transcription-coupled repair^30–32^. We also found InDels to be depleted from exonic regions of chondrocytes, but this was only significant for the non-OA control samples (p<0.0001). Significant InDel depletion in enhancer regions was observed in both non-OA control (p<0.0001) and OA lesion (p<0.0001) samples. No significant depletion nor enrichment of InDels was observed in promoter regions (Fig. 4b, Supplementary Table 6).

**Figure 4.**
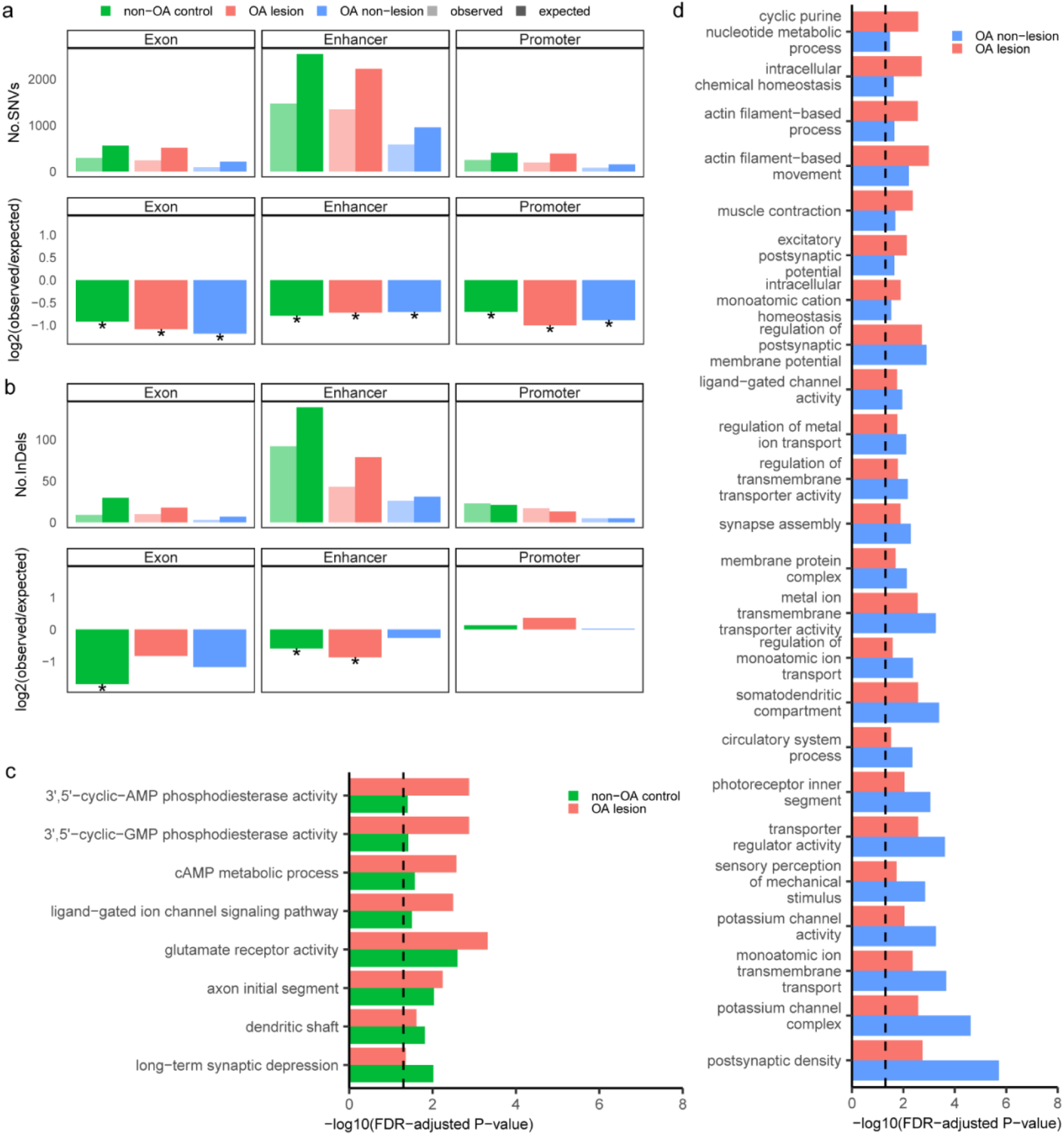
Functional enrichment analysis. **a,** Depletion of SNVs in exon, promoter and enhancer for each cartilage damage group. **b,** Depletion of InDels in exon, promoter and enhancer for each cartilage damage group. The log_2_ ratio of the number of observed and expected base substitutions indicates the effect size of the depletion in each region. Asterisks represent significant depletions per indicated disease group (P < 0.05, binomial test, one-sided). **c,** Differentially enriched GO categories (FDR-adjusted P < 0.05, permutation test) of mutated genes between OA lesion and control group. **d,** Differentially enriched GO categories (FDR-adjusted P < 0.05, permutation test) of mutated genes between OA lesion and OA non-lesion group in paired samples. x axis denotes the enrichment P value for SNVs.

We used GO (Gene Ontology) analysis to reveal potential biological processes in which the genes containing SNVs were involved. Genes containing SNVs were enriched for GO terms related to cell adhesion, cell morphology, and ion receptors in all three groups of cartilage damage status (Supplementary Table 7). The differentially enriched GO terms between the OA lesion group and the non-OA control group were mainly related to cyclic adenosine monophosphate (cAMP) metabolic process and glutamate receptor activity (Fig. 4c, Supplementary Table 7). cAMP is a secondary messenger that can influence various cellular processes; it is involved in the DNA damage response (DDR) and can either stimulate or inhibit apoptosis^33,34^. Levels of cAMP in chondrocytes play a role in controlling catabolic activity and increased cAMP levels in chondrocytes have been found to inhibit cartilage degradation^35^, suggesting that somatic mutations may affect OA progression by influencing chondrocyte metabolism and apoptosis. Glutamate receptor NMDAR regulates calcium flux into OA chondrocytes and affects the circadian clock and cell phenotype in osteoarthritic chondrocytes^36,37^. In the paired OA samples, ion transmembrane transport related GO terms were shown to be differentially enriched in the OA lesion and non-lesion group (Fig. 4d; Supplementary Table 7). Ion channel changes play a role in the pathogenesis of OA, with alteration in chondrocyte ion channels, especially calcium signaling channels, will affect extracellular matrix metabolism and regulate the pain neurotransmitters between the cartilage and the central nervous system ^38–40^. Our observation that SNVs and InDels are enriched in genes participating in GO terms related to ion transport and cAMP metabolic processes point toward the possibility that the accumulation of somatic mutations in functional genes of chondrocyte plays a role in the onset and progression of OA.

### Chondrocyte loss and apoptosis in OA lesion cartilage

Considering that the possible presence of chondrocyte apoptosis in OA cartilage may lead to elimination of chondrocytes or abnormal cell morphology, thereby resulting in bias during the isolation of seemingly healthy single chondrocytes, we analyzed morphology and apoptosis of chondrocytes in cartilage tissue sections. We checked the morphology of cartilage samples by H&E staining and observed enlarged, empty lacunae and lower chondrocyte density in OA cartilage *in situ* than in non-OA subjects (Extended data Fig. 4). Using TUNEL staining, we detected higher levels of chondrocyte apoptosis at the lesion site of OA cartilage than at the paired non-lesion site from the same patient, while apoptosis was very rare in non-OA control cartilage (Extended data Fig. 4). The empty lacunae, lower cell density and higher TUNEL signal indicate chondrocyte apoptosis in OA-lesion cartilage. Hence, it is conceivable that we missed damaged chondrocytes during single cell isolation, which could explain the lower somatic mutation burden in the OA lesions.

## Discussion

Osteoarthritis (OA) is a prevalent degenerative joint disease, with advanced age as the primary risk factor for its incidence and development^41^. It has been well recognized that aging of individuals is accompanied with genome instability, including DNA damage and somatic mutations^42,43^. In chondrocytes, DNA damage level increases with age, and OA cartilage has been reported to have significantly higher DNA damage level than non-OA cartilage^18,44,45^. This may be associated with changes in the activity of DNA damage repair, which in turn may be a mechanism to delay the progression of osteoarthritis^19,46^. DNA damage has been proposed as a universal cause of aging, which is in keeping with recent evidence that the accumulation of somatic mutations with age is a universal feature in human tissues and cell types^42,43,47,48^. The landscape of somatic mutations during aging has been well characterized in multiple organs and tissues of homo sapiens, both in normal tissues and in tissues involved in diseases, including cancer and degenerative diseases^4,24,48^. Yet, the somatic mutational landscape in chondrocytes as a function of aging and disease has been absent. Our present results now fill this gap in our knowledge and show that somatic mutations also increase with age in human chondrocytes. While OA has been found to associated with chromosomal aberrations^16^, a possible role of the vast bulk of genome instability in the form of somatic SNVs and InDels was unknown. Surprisingly, here we provide evidence that the frequency of such mutations is lower rather than higher in chondrocytes of OA lesions^17^.

The overall somatic mutation burden in human chondrocytes appears to be in the range of the other human cell types thus far analyzed, i.e., lymphocytes, hepatocytes, bronchial epithelial cells, neurons and cardiomyocytes^48^. The rate of mutation accumulation observed in chondrocytes of healthy subjects is on the high side, i.e., higher than in lymphocytes^9^ or bronchial epithelial cells^28^, but in the same range as neurons^2^ or hepatocytes^8^. Based on observations in neurons of Alzheimer’s Disease^4^, as well as the well-documented inflammation in OA lesions, we expected to see increased age-related mutation accumulation in chondrocytes isolated from these lesions as compared to normal chondrocytes. Interestingly, we found the exact opposite, i.e., a slower rate of mutation accumulation. This was significant for both SNVs and small InDels. The lack of an accelerated age-related mutation accumulation in OA lesions appeared to extend to chondrocytes from unaffected cartilage from the same OA patients. There was no difference in mutation accumulation rate between affected and non-affected cartilage from the same OA patient.

Inflammation and mechanical stresses are known to induce DNA damage through reactive oxygen and nitrogen species and there is evidence for excess DNA damage in OA chondrocytes^49,50^. We would have expected that this increase in DNA damage leads to increased DNA mutations through errors during replication and repair. However, based on our current results one could propose the alternative scenario in which the increased DNA damage causes cell death through the DNA damage response. Hence, we would retrieve only cells that have been spared excessive DNA damage and not undergone mutagenesis. Our observation that somatic mutation load in chondrocytes from superficially normal, adjacent cartilage tissue is similarly low as that in the same cells from OA lesions suggests that excessive DNA damage production and repair has already progressed. In adult articular cartilage, cell loss increases during normal aging^51^ and in an accelerated manner in OA^52^. Increased chondrocyte apoptosis is one of the hallmarks of cartilage degeneration in OA at a rate that is much higher than in normal cartilege^53,54^. Increased apoptosis due to increased DNA damage, which has been reported for OA cartilage^55^ and was observed in our present study too, could explain our observation that somatic mutation burden is lower rather than higher in OA cartilage. There is evidence that high toxicity leads to a reduced somatic mutation frequency^56^, while in tobacco smoke-induced mutations in the human lung the median SNVs frequency was also lower rather than higher in heavy smokers than that in moderately smoking individuals^28^.

Mutational signature analysis reveals that the aging-related clock-like mutation signature SBS5 accumulated in all detected chondrocytes at a frequency and accumulation rate that was independent from cartilage damage status. We did not detect another clock-like signature, i.e., SBS1, in chondrocytes, which is caused by G:T mismatches in double stranded DNA due to deamination of 5-methylcytosine to thymine prior to DNA replication, therefore is regarded as cell division/mitotic clock. The constant presence of DNA damage repair associated signature B across all three disease groups also suggests that DNA damage repair processes are the source of somatic mutations in chondrocyte.

We also extracted *de novo* signatures for both SNVs and InDels that showed a significant difference between OA and non-OA control samples, specifically identified as signature C and signature 1, which are associated with defective NER and MMR. An age-related decline in DNA damage repair has been reported in chondrocytes^20^. In addition, during aging and after trauma, the physiological hypoxic environment in which chondrocytes reside is compromised^57^. Moreover, chondrocytes are subjected to higher mechanical stress, which could cause higher level of DNA damage^50,58^. Increased damage and reduced repair led to the result of accumulated DNA breaks in chondrocytes from OA and aged cartilage. However, DNA damage repair processes are not perfectly precise, errors in DNA damage repair generate mutations. Therefore, in OA chondrocyte, where DNA damage repair is reduced, alongside a higher DNA break burden, a relatively lower burden of somatic mutation was observed.

Through Gene Ontology (GO) analysis, we observed differential enrichment of somatic mutations in gene terms important for chondrocyte functions and cartilage homeostasis between OA chondrocytes and controls. For example, in OA chondrocytes, mutations were enriched in terms related to cAMP metabolism and ion transportation. cAMP is essential in a wide variety of cellular pathways, including the DNA damage response and the regulation of apoptosis. In chondrocytes the cAMP level is important in regulating the inhibition of cartilage degradation. Ion transport in chondrocytes can regulate extracellular matrix metabolism and OA-related pain. Somatic mutations may influence OA incidence and progression by affecting chondrocyte metabolism and apoptosis. The observed differentially enriched GO terms in OA and control chondrocytes are consistent with the possibility that somatic mutation accumulation has adverse effects at both cellular and systemic levels.

In summary, the results presented offer insight for the first time into the somatic mutational landscape of chondrocytes, the cell type that is central in the etiology of osteoarthritis, a major age-related disease. Similar to what we and others have recently shown for other human cell types, somatic mutations accumulate with age in human chondrocytes as well. However, in contrast to expectations, we show that the well-documented increased inflammation of OA lesions does not result in increased somatic mutation burden of chondrocytes isolated from these lesions. Instead, we observed a slower age-related mutation accumulation in OA chondrocytes than in the same cells isolated from control subjects. The most likely explanation for this reduced rather than increased somatic mutation accumulation rate is inflammation, which is highly toxic, inducing ample DNA damage in the chondrocytes in OA lesions. We suggest that this increased DNA damage leads to excess cell death. Increased apoptosis is well-documented in OA lesions, which our own data fully confirm. In this respect, it would be of interest to study somatic mutations in degenerative tissues of other diseases, most notably atherosclerotic plaques, which have also been associated with increased inflammation^59^.

Finally, one limitation of our present study is the lack of insight into genome structural alterations as part of the somatic mutational landscape. Such mutations are rare but much more impactful than SNVs or INDELS but cannot be detected at the somatic level using current methods, which themselves have become available only recently. Nevertheless, we cannot rule out the possibility that increased inflammation preferentially induces this type of mutations.

## Methods

### Data reporting

No statistical methods were used to predetermine sample size. The experiments were not randomized, and the investigators were not blinded to allocation during experiments and outcome assessment.

### Human tissue samples and selection of cases of participants

Cartilage samples of hip femoral heads of 17 individuals, and peripheral blood of 4 donors were obtained from the Department of Orthopedics, Shanghai Sixth People’s Hospital, Shanghai Jiao Tong University School of Medicine. Tissue collection and distribution for research and publication was conducted according to protocols approved by the Independent Ethics Committee of Shanghai Sixth People’s Hospital (approval numbers: 2016-KY-001(K)-(5), 2021-122), and after provision of written authorization and informed consent. Hip articular cartilage samples were collected from patients with osteoarthritis undergoing hip/knee replacement surgery; and patients undergoing hip/knee replacement surgery for other reasons.

### Cartilage dissociation

Cartilage samples collected from surgery were kept in pre-cooled sterile D-PBS (pH 7.4, Beyotime C0221D) and transferred to a biosafety cabinet on ice. Samples were then washed 3 times by rinsing with pre-cooled D-PBS and cut into small pieces of about 1mm^3^ using a sterile surgery blade. Cartilage pieces were incubated in 0.25% trypsin-EDTA (∼500µl/100mg cartilage, GIBCO 25200056) at 37℃ 5% CO_2_ for 30 min while mixing on a rotator at 25-30 rpm. Trypsin was then removed and tissue pieces were washed once with DMEM (GIBCO 11965-092). Cartilage pieces were then further dissociated in 4mg/ml collagenase II in DMEM (∼500µl/100mg cartilage, Sigma-Aldrich C1764) at 37℃, 5% CO_2_ for 1.5-2 hours, while mixing on a rotator at 25-30 rpm. The suspension was then filtered through a 70-micron cell strainer (Falcon 352350) and the cell suspension collected by centrifugation at 300g for 5min at room temperature; the cell pellet was then washed once with DMEM. Cells were counted and checked for viability by staining with trypan blue (Beyotime ST798) with Countess II FL Automated Cell Counter (Invitrogen AMQAF1000). Nuclear integrity was checked using PI (Beyotime ST511) and DAPI (Beyotime C1002) staining on an inverted fluorescent microscope (NIKON Ti2-FL).

### Isolation of single chondrocytes

Single cells were obtained using a cellenONE X1 (Cellenion) automatic single cell sorting system, from a suspension of 2^5^ cells/ml, filtered through a 40-micron cell strainer (pluriSelect 43-10040-60). Following instrument operation instructions, single cells were collected with a size between 10 and 20 microns, elongation below 1.8, and circularity between 1.0 and 1.1 into 96 well PCR plates (Axygen PCR-96-FS-C) or PCR strip tubes (Axygen PCR-0208-C) containing 2.5μl PBS, snap-frozen on dry ice, kept in -80℃ until further use.

We also isolated single chondrocytes using the manual single-cell isolating platform QIAscout (Qiagen). For QIAscout, we seeded 3000∼6000 cells to 1 QIAscout array raft (Qiagen 928031), collected individual cells into 0.2-ml PCR tubes containing 2.5μl PBS; they were snap-frozen on dry ice and kept at −80 °C until further use.

### Single-cell whole-genome amplification

Whole genomes of single chondrocytes were amplified using a single-cell multiple displacement procedure as described previously^7,21^. 1 ng human genomic DNA and DNA-free PBS solution were applied to SCMDA in parallel with single cells as positive and negative controls, respectively. Amplification products were purified with AMPureXP beads (Beckman Coulter A63881). Product DNA concentrations were assessed using a Qubit 1X dsDNA HS Assay Kit (Invitrogen Q33231). Product size was determined by agarose gel electrophoresis, with a size of over 10 kb considered acceptable. Amplification uniformity was checked using real-time PCR (PowerUp SYBR Green Master Mix, Applied Biosystems A25778) of 8 separate loci ^7,60^. Samples with at least six out of eight loci positive were considered acceptable and further subjected to library preparation and WGS.

### Genomic DNA extraction

Human bulk genomic DNA was extracted from 0.2 million chondrocytes collected from the same cartilage sample after single cell isolation using the QIAamp DNA Mini Kit (Qiagen 51304) according to the manufacturer’s protocol. DNA concentration was quantified using Qubit 1X dsDNA HS Assay Kit (Invitrogen Q33231).

### Library preparation and WGS

100∼500ng genomic DNA or amplified single chondrocyte genomic DNA were used as input to prepare sequencing libraries for the Illumina platform with NEBNext Ultra II FS DNA Library Prep Kit (NEB E7805). Briefly, genomic DNA was incubated with fragmentation enzyme for 8 min to generate fragments in size between 300∼400bp. After ligation of adaptor and beads purification for size selection, libraries were enriched by a 5-round PCR. The libraries were sequenced on Illumina NovaSeq 6000 S4 sequencing platform to 30X depth using a 2×150 paired-end mode.

### Alignment for whole-genome sequencing

WGS data were processed with the Sarek pipeline implemented within nextflow-core (nextflow version 19.10). The raw sequencing reads were trimmed to remove adapter and low-quality nucleotides by Trim Galore (version 0.6.4). The trimmed reads were aligned to the human reference genome (GRCh37 with decoy) using BWA (mem; version 0.7.13). PCR duplications were removed by samtools (rmdup; version 1.9). To correct mapping errors made by genome aligners, the known InDels and SNPs were collected from the 1000 Genomes Project (phase I) and dbSNP (build 138). Then InDels realignment and base quality score recalibration was performed based on known InDels and SNPs via Genome Analysis Toolkit (GATK, version 4.1.7).

### Calling germline variants from bulk sequencing

Germline SNVs and small InDels were called by HaplotypeCaller. The variants with GATK quality score ≥ 30 were maintained and further filtered based on the recommendation of GATK as ‘QD < 2.0, FS > 60.0, MQ < 40.0, MQRankSum < -12.5, ReadPosRankSum < -8.0, 20 SOR > 3.0’ for SNV and ‘QD < 2.0, FS > 200.0, ReadPosRankSum < -20.0, SOR > 10.0’ for InDel.

### Calling somatic small variants

Somatic mutations between each single cell and the corresponding bulk were identified by SCcaller (version 2.0.0)^7^. Only mutations on autosomes were considered. Known heterozygous SNPs were called from bulk DNA using HaplotypeCaller. To obtain high-quality mutation calls, we only considered heterozygous SNPs with position coverage ≥ 20X, GATK phred-scaled quality score ≥ 30 and dbSNP annotations. Somatic mutations with a depth less than 20X or those supported by bulk data were filtered out by default. For InDels, we required 30X depth and phred-scaled quality score ≥ 25 to maintain high accuracy results. Mutations overlapping with known SNPs in dbSNP were also annotated and removed using SnpEff (version 4.3).

### Variants annotation

All identified somatic mutations were annotated according to the gene definitions of GRCh37. Annotations from ANNOVAR were used to identify mutations falling in the following positions: intergenic, upstream (within 1 kb region upstream of transcription start site), 5′ UTR, exonic (coding sequence, not including untranslated regions), 3′ UTR, downstream (within 1 kb region downstream of transcription start site), splicing (within intronic 2 bp of a splicing junction), intronic. The regulatory features in Ensembl were used to identify mutations fall in following features: promoters, enhancers, open chromatin regions, transcription factor binding, CTCF binding sites. All annotation were performed using ANNOVAR.

### Estimating mutation frequencies

The frequency of somatic SNVs/InDels per cell was estimated with the normalization of genomic coverage and sensitivity:

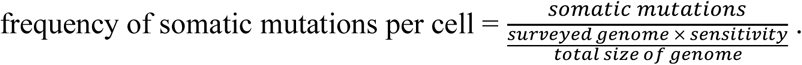

The surveyed genome per single cell was calculated as the number of nucleotides with mapping quality ≥40 and coverage depth ≥20X for SNVs and ≥30X for InDels, which are also found in its corresponding bulk DNA. The surveyed genome for each single cell divided by the total size of the genome is equivalent to their sequencing coverage. The sensitivity of somatic mutation calling in the single cell was estimated as the ratio of the number of heterozygous SNPs detected in single cell to the total number of heterozygous SNPs detected in its corresponding bulk DNA, with sequencing depth requirements of at least 20X for bulk DNA, 20X for single-cell SNVs, and 30X for single-cell InDels. For InDels, we also required a quality score of SNPs in single cells ≥25. Only samples with sensitivity ≥20% and coverage ≥15% at 20X sequencing depth were considered qualified sequencing samples and included in all subsequent analyses.

### Statistical methods

To estimate the effects of age on somatic mutation burden and on the somatic mutations attributed to specific signatures, we fitted somatic mutation number of each single cell with an LME model. In the LME model, the response variable was the frequency of somatic mutations per cell. Disease status and age were modelled as fixed effects, and sample groups were modelled as random effects, because chondrocytes from a donor may be correlated owing to shared biological environment.

To estimate the effects of age on somatic mutation burden and somatic mutation burden attributed to specific signatures in the overall samples, we fitted the model:

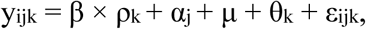

where y_ijk_ is the somatic mutation frequency in chondrocytes i from damage status group j of donor k, β is the fixed-effect of age, ρ_k_ is the age of donor k, α_j_ is the fixed-effect of disease status, μ is the number of somatic mutation frequency at birth, θ_k_ is the random effect, and ε_ijk_ is the measurement error.

To estimate the effects of age on somatic mutation burden and somatic mutation burden attributed to specific signatures within damage status groups, we fitted the model:

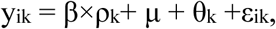

where y_ik_ is the somatic mutation frequency in chondrocytes i from sample k, β is the fixed effect of age, ρ_k_ is the age of donor k, μ is the number of somatic mutation frequency at birth, θ_k_ is the random effect, and ε_ik_ is the measurement error.

The mutation accumulation rate in each group was calculated by dividing mutation frequencies by age. To test the difference in the accumulation rate between different damage status groups, we performed a tow sided t-test.

In all models, the statistical significance was assessed using likelihood ratio tests. All analysis were conducted with R, Linear mixed effect model was estimate using function lmer of R package lme4 (v.1.1-23) and lmerTest (v.3.1-2) R packages.

### Mutational signatures

All identified mutations were pooled into three disease groups, *de novo* mutational signatures were extracted by the NMF-based mutational signature framework using MutationalPatterns (v.3.4.1)^22^.

We calculated the frequency of SNVs in the 96-trinucleotide contexts and extracted three *de novo* mutation signatures from 96-trinucleotide contexts of SNVs. For InDels, two *de novo* mutation signatures were extracted from their ID83 channel. To identify the potential origin of the mutational spectra, *de novo* signatures were compared with cancer mutation signatures from the COSMIC database (version 3.2) by calculating cosine similarity. In addition, we performed signature refitting to quantify the contribution of COSMIC signatures to the mutational profile of three sample groups, and selected the signatures whose contribution is greater than 5% as the new signature set for signature refitting.

To analyze the relationship between mutational signatures and age and disease status, we calculated the adjusted number of somatic mutations contributing to a specific mutational signature in each single cell, and fitted a generalized mixed-effect model.

### Genomic distribution analysis

To test whether the somatic mutations appear more or less frequently than expected in genes, promoters, promoter-flanking, and enhancer regions, we loaded the UCSC Known Genes tables as TxDb objects and the regulatory features for hg19 from Ensembl using biomaRt. We tested for enrichment or depletion of somatic mutations in the genomic regions per disease group using a one-sided binomial test with MutationalPatterns, which corrects for the surveyed genomic areas.

### Functional enrichment analysis

Functional enrichment analysis was performed using the R package GOseq (v.1.34.1). We took RefSeq genes as background genes and built the SNV-gene table according to whether any SNV was present in RefSeq genes in any single cell for each group. For each RefSeq gene, we assigned a binary value ‘0’ or ‘1’ according to whether any SNV is located in the corresponding gene. We performed GO analysis. Genes without any GO annotation was ignored when calculating the total gene counts. A probability weighting function in GOseq was applied to control for potential gene length bias. The Wallenius approximation method was used to test the enrichment of SNVs, and the false discovery rate (FDR) method was further applied for the correction of multiple hypothesis testing. GO terms with fewer than 10 member genes with SNVs were excluded to avoid ascertainment bias. GO terms with more than 1,000 member genes were also excluded.

To identify GO terms that were specifically enriched in a certain group, we performed a permutation test among all GO terms with P < 0.01. For each permutated GO term, we compared the observed rank difference in GOseq’s P between one group and its corresponding control group against the expected null distribution, which was estimated by 1,000 rounds of random shuffling of the SNV-gene tables. The FDR method was applied for correcting multiple hypothesis testing.

To test whether the difference between mutation frequency of OA high-confidence effector genes were statistically significant, we conducted a permutation test with resampling. We randomly selected a list of genes equal in size to the OA high-confidence effector gene set and calculated the mutation frequency. We repeated the resampling process 1000 times and compared the obtained results to the mutation frequency in OA high-confidence effector genes.

### In situ detection of apoptosis and proliferation

Cartilage samples were cut into rectangle pieces of 0.1-0.2 cm^2^ and fixed in 4% paraformaldehyde followed by paraffin embedding. Cartilage sections were then dewaxed and treated following the instruction from manufacturer of H&E staining kit (Beyotime C0105S). Cartilage sections were stained with H&E staining kit and apoptotic chondrocytes in cartilage samples were stained using terminal deoxynucleotidyl transferase mediated dUTP nick end labelling (TUNEL) method (VAZYME A112), the 3’-hydroxyl (3’-OH) end of DNA fragments were labeled with FITC-12-dUTP. FITC-12-dUTP labeled DNA and DAPI labeled nuclei were observed by fluorescence microscopy.

### Data availability

Somatic-mutation calls, including single-base substitutions and InDels from all 100 samples were included in the supplementary materials.

### Code availability

Custom codes for statistical analysis, mutational signature analysis, permutation analysis, are available through GitHub (https://github.com/zhengchen98/OA-WGS-analysis).

## Acknowledgements

We thank the CSCOmics Single-Cell Isolation & Handling Core, Single-Cell Genomics Core and Single-Cell Bioinformatics Core of CSCOmics for their assistance and helping in carrying out and completing this project. This study was financially supported by the National Natural Science Foundation of China grant (82172461, 82002339, 81820108020), Shanghai Frontiers Science Center of Degeneration and Regeneration in Skeletal System (BJ1-9000-22-4002), Shanghai Municipal Hospital Orthopedic Specialist Alliance, Shanghai Municipal Health Commission key priority discipline project; Shanghai Spinal Disease and Trauma Orthopedics Research Center (2022ZZ01014).

## Author contributions

J.V., C.Q.Z., J.J.G. and X.D. conceived and supervised the study. J.V., P.J.R., M.S.L. and A.Y.M. designed the experiments. C.Q.Z., J.J.G., Y.D.P. and N.N.L. provided clinical procedure and specimen-specific study expertise. X.D., C.Z., S.X.S., and W.X.X. set up the analysis workflow. J.J.G., Y.D.P., and Q.Y.W collected samples from joint replacement surgeries. P.J.R., C.Z., Y.Q., Z.Z.L. and M.Z. performed the experiments. J.H. provided sequencing support. C.Z., P.J.R., X.D. and J.V. analyzed the data. J.V., P.J.R., C.Z., and J.J.G., wrote the manuscript.

---

**Extended data Figure 1.**
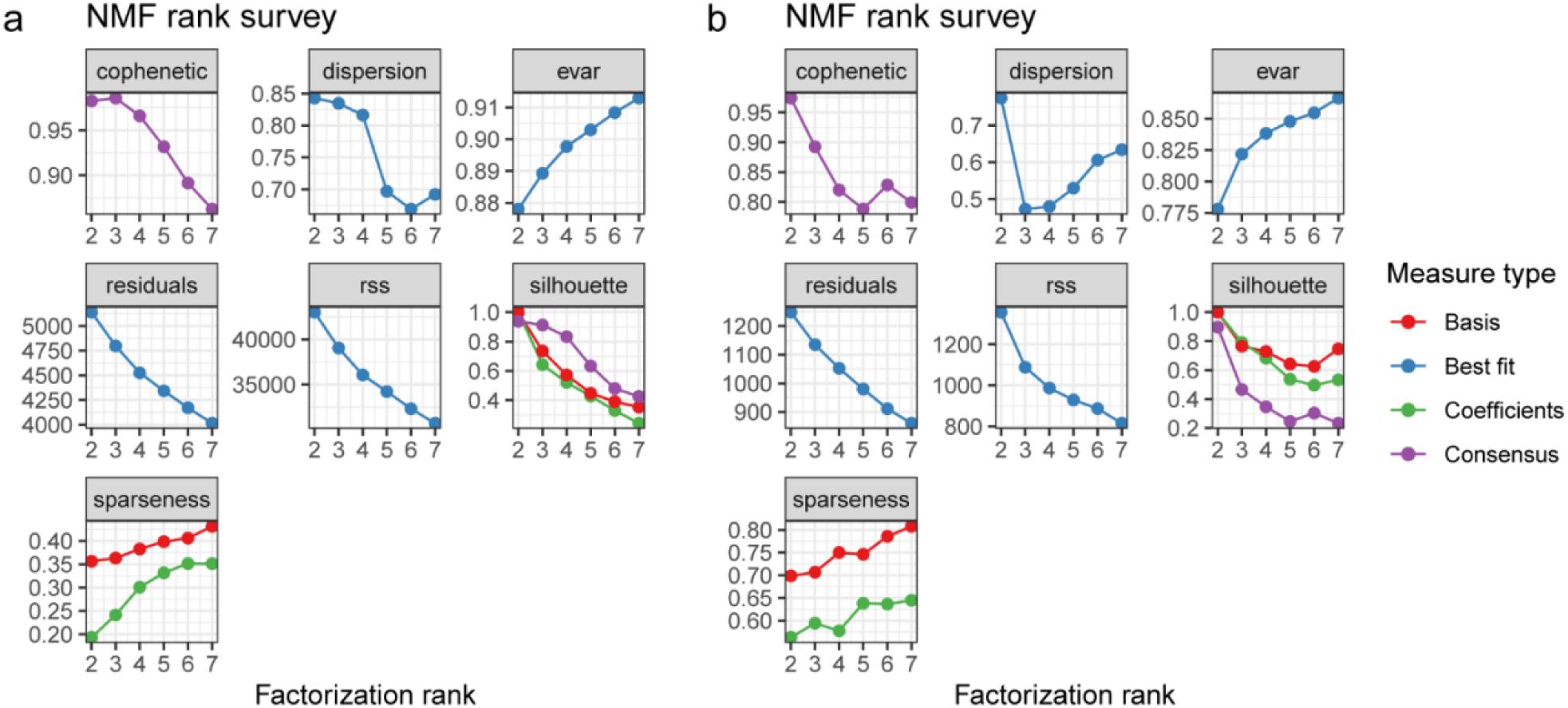
Signature metrics for *de novo* mutational signature analysis. *De novo* mutational signature analysis was performed using nonnegative matrix factorization (NMF), in which the factorization rank is critical to define the number of signatures used to decompose the target matrix of SNVs. **a,** 3 signatures can maximize the cophenetic and best fit the observed SNV matrix. **b,** 2 signatures can maximize the cophenetic and best fit the observed InDel matrix.

**Extended data Figure 2.**
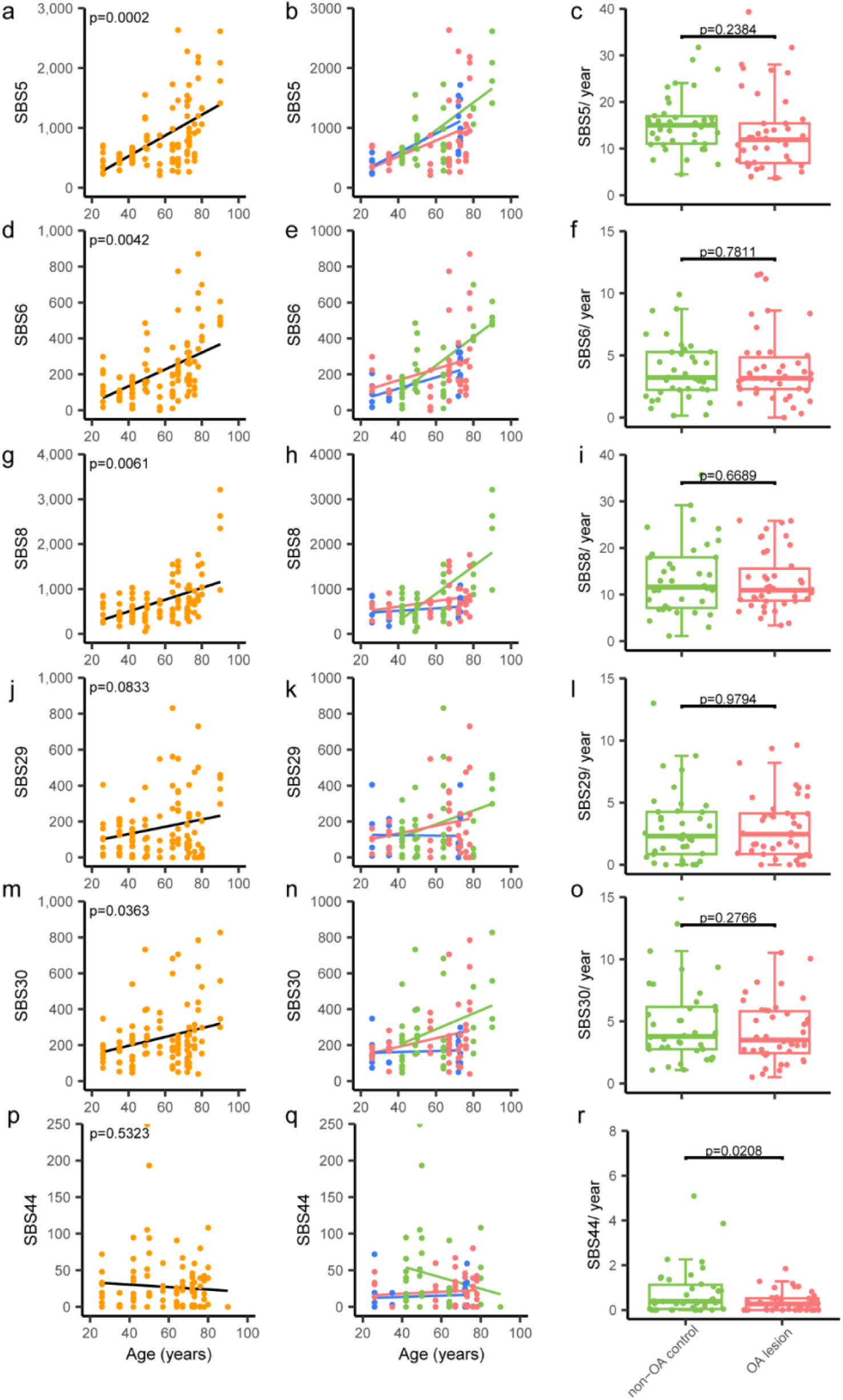
Contributions of mutational processes to chondrocytes. **a, d, g, j, m, p,** Contribution of COSMIC signatures to SNVs versus age, with linear mixed-effect regression lines. Each data point indicates the number of mutations contributing to the signature per single chondrocyte (n=100). P values were obtained by linear mixed-effect models. **b, e, h, k, n, q,** Contribution of COSMIC signatures to SNVs versus age in three damage status groups, with linear mixed-effect regression lines. Each data point indicates the mutation frequency per single chondrocyte. **c, f, i, l, o, r,** The per year accumulation rate of COSMIC signatures contributing to the SNVs in different damage status groups, with boxes indicating median values and interquartile ranges of the non-OA control and OA lesion, respectively. Each data point indicates the per year mutation accumulation rate of each single chondrocyte contributing to the signature.

**Extended data Figure 3.**
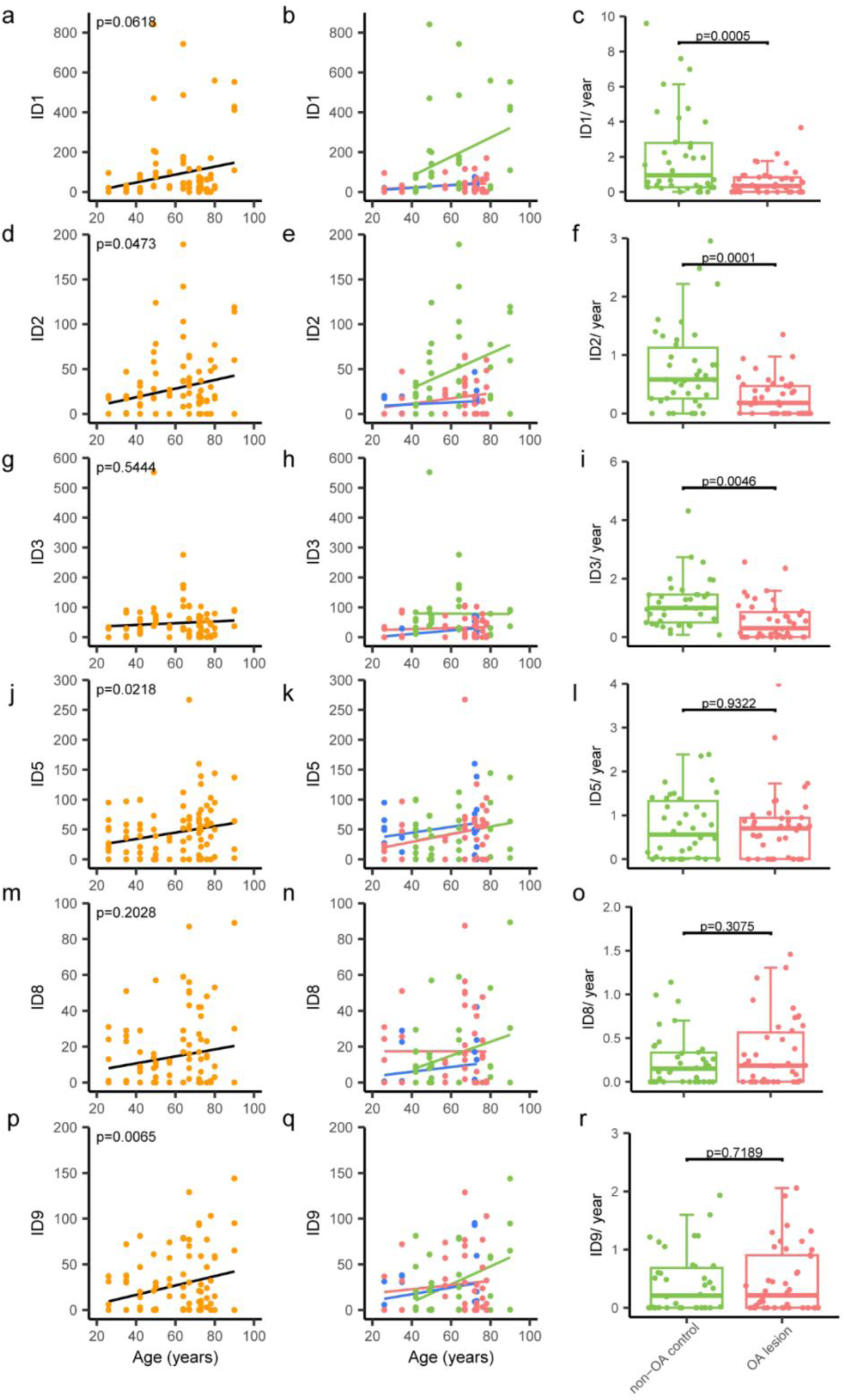
Contributions of mutational processes to chondrocytes. **a, d, g, j, m, p,** Contribution of COSMIC signatures to InDels versus age, with linear mixed-effect regression lines. Each data point indicates the number of mutations contributing to the signature per single chondrocyte (n=100). P values were obtained by linear mixed-effect models. **b, e, h, k, n, q,** Contribution of COSMIC signatures to InDels versus age in three damage status groups, with linear mixed-effect regression lines. Each data point indicates the mutation frequency per single chondrocyte. **c, f, i, l, o, r,** The per year accumulation rate of COSMIC signatures contributing to InDels in different damage status groups, with boxes indicating median values and interquartile ranges of the non-OA control and OA lesion, respectively. Each data point indicates the per year mutation accumulation rate of each single chondrocyte contributing to the signature.

**Extended data Figure 4.**
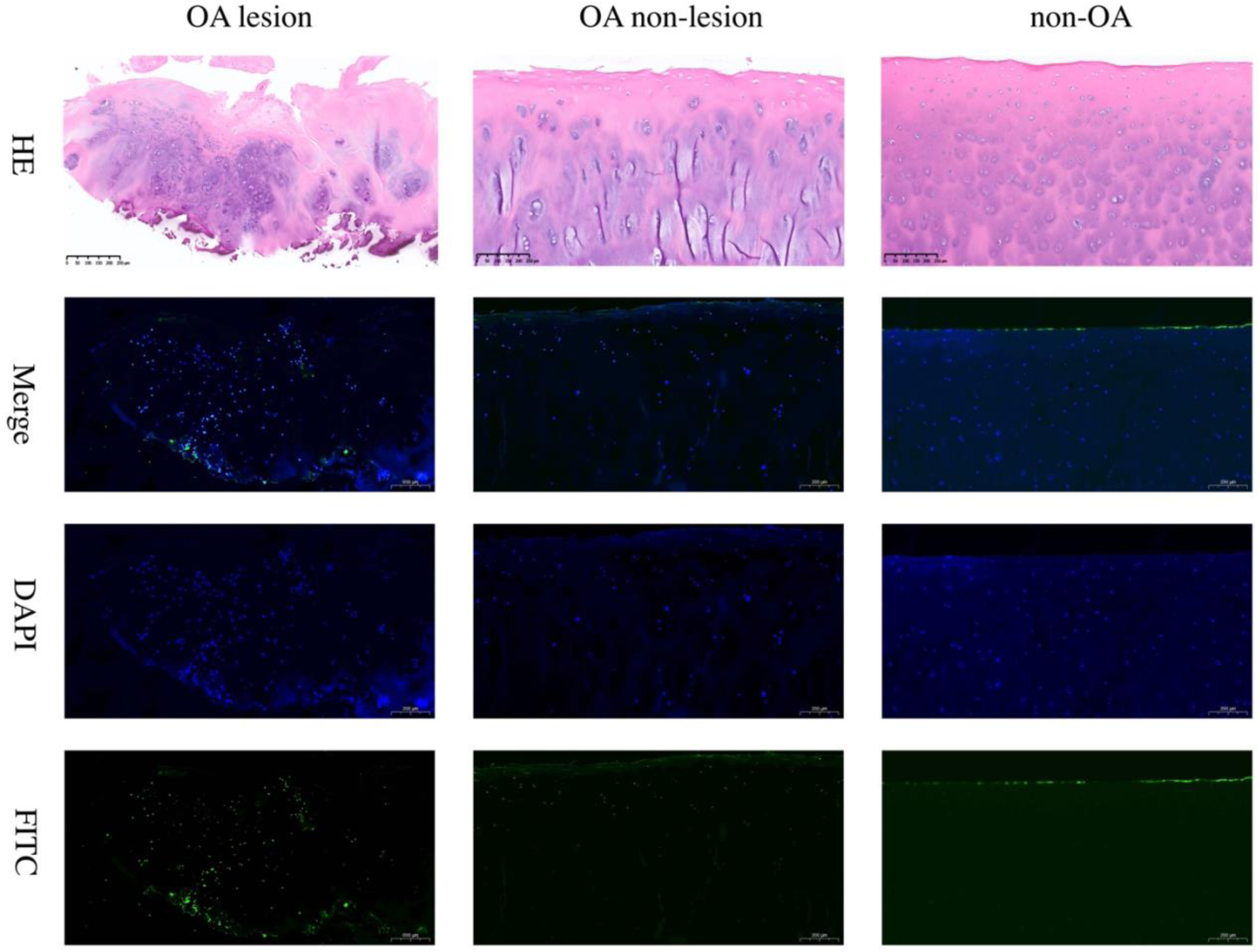
H&E and TUNEL staining of femoral head cartilage sections. The representative images of H&E staining (the upper row) and TUNEL staining of apoptosis (the lower 3 rows) of lesion (left) and non-lesion (middle) sites of OA femoral head cartilage, and non-OA femoral head cartilage (right), respectively.

